# DotMotif: An open-source tool for connectome subgraph isomorphism search and graph queries

**DOI:** 10.1101/2020.06.08.140533

**Authors:** Jordan K. Matelsky, Elizabeth P. Reilly, Erik C. Johnson, Jennifer Stiso, Danielle S. Bassett, Brock A. Wester, William Gray-Roncal

## Abstract

Recent advances in neuroscience have enabled the exploration of brain structure at the level of individual synaptic connections. These connectomics datasets continue to grow in size and complexity; methods to search for and identify interesting graph patterns offer a promising approach to quickly reduce data dimensionality and enable discovery. These graphs are often too large to be analyzed manually, presenting significant barriers to searching for structure and testing hypotheses. We combine graph database and analysis libraries with an easy-to-use neuroscience grammar suitable for rapidly constructing queries and searching for subgraphs and patterns of interest. Our approach abstracts many of the computer science and graph theory challenges associated with nanoscale brain network analysis and allows scientists to quickly conduct research at scale. We demonstrate the utility of these tools by searching for motifs on simulated data and real public connectomics datasets, and we share simple and complex structures relevant to the neuroscience community. We contextualize our findings and provide case studies and software to motivate future neuroscience exploration.

## 1 Introduction

Modern nanoscale connectomics research commonly involves the conversion of microscopy imagery data into a graph representation of connectivity, where nodes represent neurons, and directed edges represent the synapses between them [1]. This process enables researchers to convert terabytes or even petabytes of imagery into megabytes or gigabytes of graph data. Conversion to a network format reduces the cost and complexity of interrogating the data, at the expense of losing information about cellular morphology [2, 3]. Though this graph representation uses substantially less storage-space on disk, answering even seemingly simple network questions (e.g., identifying local graph structure around a particular neuron, or comparing the downstream targets of a certain cell type) may still exceed the computational power, timelines, and budgets available to many research teams, due to the exponential nature of many graph algorithms [4, 5]. This issue is often ameliorated by including node or edge attribute constraints in the search, though this places an additional limitation on the types of questions that a researcher can expect to ask of a connectome dataset.

Connectomics researchers have begun to address these challenges of large-scale graph analysis by adopting existing large-scale graph management software from other domains, such as graph databases, and by enforcing consistent, well-architected data schemas [6, 7]. These systems provide performant and cost-effective ways to manipulate larger-than-memory graphs, but tend to require familiarity with complex and nuanced graph query programming languages such as Gremlin or Cypher. Though graph databases continue to grow in popularity, the expertise to administer or use these technologies is still not common in the biological sciences.

In order to make the study of connectomes accessible and computationally efficient, we developed *DotMotif*, an intuitive but powerful graph tool designed to reduce the expertise and time required to begin interrogating biological graphs of any size. DotMotif acts as an interface to common graph management systems such as the NetworkX Python library or the Neo4j graph database, abstracting the intricacies of subgraph-query design and enabling researchers to focus on field-specific scientific inquiry. DotMotif assists researchers in query design by exposing an intuitive, simple syntax that automatically validates and optimizes user queries, and adapts to different graph tool backends without the need for additional user input. We present the DotMotif Python package and software architecture, which enables researchers to write queries while remaining agnostic to underlying technologies. We demonstrate DotMotif’s utility by assessing its advantages over manual query design, and we share several use-cases inspired by ongoing research in the connectomics community. We also share examples of how these findings might inform future work, including efforts to design better local connectivity rule sets, establish expected motif prevalence, and produce better generative models of a connectome for evaluation, null model design, or for comparison across species, brain region, and modality.

## 2 Background

Many of the fundamental questions of modern neuroscience rely on the study of simple circuits of neural connectivity in the brain. These simple circuits are hypothesized to be repeated and reused many times to perform a similar role. Though they might comprise only a small number of neurons, as shown in **Figure 1**, these simple circuits or graph *motifs* may be critical to understanding the functional role of larger structures in the brain, such as cortical columns or other modules of computational importance [8, 9, 10]. Due to the nature of available high-resolution imaging modalities such as electron microscopy, it is not always feasible to collect functional as well as anatomical data at synaptic resolution. In this work, we define motifs as repeating subgraphs, but do not make assumptions about their computational significance.

**Figure 1:**
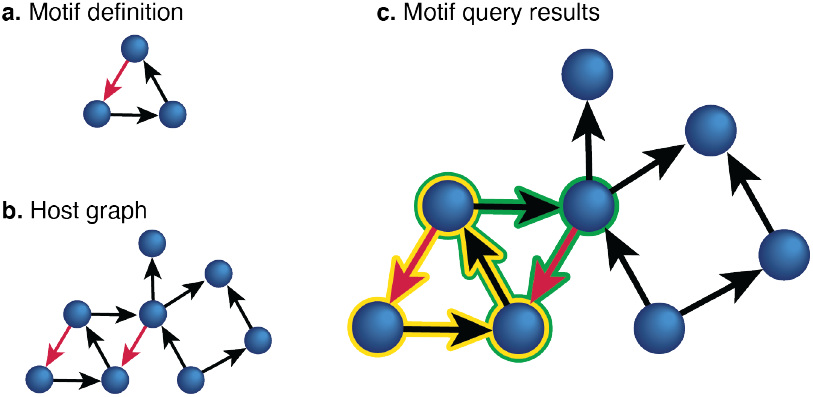
Subgraph search. Subgraph search is formally defined in the *Background* section. **a.** A motif query is defined. In this example, the query is a simple directed triangle. One edge (red) has been assigned an attribute constraint (perhaps a neurotransmitter type or weight threshold). **b.** A directed search graph, or “host” graph. Two edges in the host graph share the same attribute as the constrained edge in the motif. **c.** The motif is discovered in the search graph. Two unique detections are shown here, highlighted in yellow and green. Note that edges and nodes in the host graph may appear in more than one mapping.

Identifying these motifs is one motivation for the field of *connectomics*, the study of the brain through the lens of its connectivity. Many related research efforts seek to construct graph representations of the brain, commonly represented by the notation *G* = (*V, E, A*), where a graph *G* is made up of nodes *V*, representing neurons; (optionally directed) edges *E*, representing synapses between neurons; and arbitrary attributes *A*, which may be associated with vertices, edges, or the graph as a whole. For example, node attributes might include qualities such as cell type or functional information. Edge attributes might include synapse weight or neurotransmitter type. Graph attributes might include the species or individual from which the connectome was generated. Neuroscience questions may be reformulated as analyses on a graph, and the neuroscientist may add graph theory to the toolbox of strategies with which to understand the brain [8, 11]. Such questions include searching for specific subgraph structures, investigating the connectivity of specific neuron cell types or categories, proofreading connectomes for accuracy, and generating summary statistics on the graph as a whole [12, 13, 3, 14, 15, 16, 10].

Many graphs generated by the connectomics community in recent years have spanned multiple gigabytes of hard drive space [2, 10], rendering conventional graph toolkits, such as the common NetworkX library [17] (or its counterparts in other programming languages) under-powered to address the needs of the scientific community. These tools, which often require all graph data to be stored in RAM, would require impractically expensive compute hardware in order to run fully in-memory, and would require impractically long timelines in order to run while swapping data from memory to disk. Instead, some teams [6] have opted to leverage “out-of-memory” tools, such as Neo4j [7], Cayley [18], or other graph databases [19, 7], which operate on graph data much like conventional relational databases operate on tabular data. Despite their power, such tools require expertise in specialized query languages such as Cypher or Gremlin, much like relational databases require knowledge of languages such as SQL. Developing or hiring for this sort of domain expertise may be impractical for many neuroscience research laboratories and is independent from the core expertise needed to formulate and test neuroscience hypotheses. The need for computationally efficient, accessible, and intuitive neural graph analysis tools motivates this work.

Motif-based connectomics research is an emerging neuroscience sub-discipline. This community currently depends upon closed-form matrix algebra subgraph-counting techniques, which are only established for very specific types of motifs (e.g. fully connected motifs or star graphs). More complicated motifs, such as those which include information about cell or synapse types, or those with more topologically interesting graph connectivity, are currently understudied largely due to this technical gap. In order to facilitate directed hypothesis testing in large-scale connectomics, the community will need motif search tools that are able to incorporate information about cell types, local morphology, functional and structural data, and other graph, node, and edge attributes.

## 3 Results

For DotMotif to be useful to the neuroscience community, it is important for potential end users to see its practical utility. We share illustrative DotMotif experiments run on a set of publicly-available, seminal connectome datasets. We first validate our results by comparing against existing online data, and then share a network-analysis result on both partial as well as complete connectomes. Lastly, we share performance benchmarks for graphs of various sizes. Collectively, these results serve to illustrate how our tools enrich the connectomics toolkit.

### 3.1 Datasets

We demonstrate the use of our tool on both near-complete (most or all of the organism’s neurons are included in the graph) connectomes as well as on partial connectomes. Here, we compare connectomes of the invertebrate nematode *C. elegans* [23, 20], the partial connectome of the invertebrate fruit fly from the Howard Hughes Medical Institute Janelia Hemibrain [2] project, and the partial vertebrate mouse visual cortex connectome from the Intelligence Advanced Research Project Activity (IARPA) Machine Intelligence from Cortical Networks (MICrONS) project [24, 25, 10]. Further dataset details are available in **Table 1**. These published graphs do not adhere to a particular database schema or storage system, and each has different node and edge attributes. Despite these differences, all three can be analyzed using the techniques we detail here.

**Table 1:** Connectome graphs reviewed in this work. The *C. elegans* connectome was derived from the July 2020 *C. elegans* adjacency matrix at *WormWiring.org*, and synapses were filtered to only include chemical synapses for this analysis [20, 21]. The MICrONS v185 graph was derived from the proofread soma subgraph at *microns-explorer.org* [10, 22]. The *Hemibrain* v1.2 dataset was accessed through the neuPrint system [6].

| Dataset | Nodes | Edges | Density | Species/Area |
| --- | --- | --- | --- | --- |
| *C. elegans* chemical synapses | 296 | 3427 | 0.064 | *Caenorhabditis elegans* |
| MICrONS v185 | 334 | 1736 | 0.031 | *Mus musculus* visual cortex |
| Hemibrain v1.2 | 21,740 | 3,550,404 | 0.011 | *Drosophila melanogaster* |

### 3.2 Tool validation: Declarative queries

As a simple first step towards tool validation, we begin by demonstrating the utility of DotMotif in performing declarative queries on a large connectome graph. In order to illustrate compatibility with data stored in the neuPrint data format [6], we ran a DotMotif search query on the partial *Drosophila melanogaster* connectome (dubbed *“Hemibrain”*) [2]. We then validated these results with the neuPrint API at *neuprint.janelia.org* [6]. In this example motif query (**Figure 6a**), we wanted to find all *Antennal Lobe* inputs to *Kenyon Cells* with weights within a certain range. We identified seven instances of this motif (centered around antennal lobe neurons with IDs 1917188956, 1825085656, 5813063239, 1887163927, 1917188956, 1825085656, and 1037293275). With these references to neurons in the host graph, we were equipped to look for further patterns among these neurons. In order to confirm that the query above returned results that were identical to those obtained from the canonical neuPrint server, we validated these results with the equivalent Cypher command (**Figure 6b**) in the neuPrint web application. This exercise demonstrates DotMotif can be used to easily convert declarative neuroscience questions into succinct, efficient graph queries.

### 3.3 Comparing undirected subgraph searches across connectomes

As an illustration of the minimal configuration requirements of the DotMotif package, we searched the human-proofread subgraph from the IARPA MICrONS project [24, 25, 10] for all undirected graphs of size |*V*| ≤ 6, with the ordering taken from the *Atlas of Graphs* [26, p. 8–30]. We then counted the number of times each subgraph appeared. For example, we counted all undirected triangles (*Atlas of Graphs* ID #7) and discovered that there were 6894 unique triangle monomorphisms in the MICrONS graph when ignoring edge direction. There were 123,264 unique undirected rectangle (*Atlas of Graphs* ID #16) monomorphisms. The only undirected graph in this set with *no* occurrences in the MICrONS graph was Motif ID #208, which was the complete graph of 6 nodes. We publish this complete dataset of all motifs and their respective counts for community use (see *Supplemental Materials*).

To contextualize these results, we calibrated the parameters of four simple random graph models (Erdős– Rényi [27], Undirected Geometric [28], Watts-Strogatz [29], and Barbarási-Albert [30]) to match the graph density 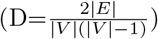 of each estimated connectome, and ran the same count of subgraph motifs on samples taken from each random graph distribution. These models were chosen for their simplicity and small parameter spaces. We similarly performed the same subgraph motif counts on the configuration model (X-swap) [31] — a degree-preserving graph randomizer used in previous neuroscience efforts [10]. Unlike the other models, this model begins with an existing connectome graph and performs arbitrary swaps of edges such that the overall in- and out-degrees of each node remains the same, but the precise pattern of connectivity changes [32]. More details about the random graphs used here are available in *Methods*. The graph densities were set to 0.03 for the MICRONS graph and 0.06 for the *C. elegans* graph (**Table 1**). In order to compare these results with a complete connectome example dataset, we ran this count-scan on all chemical synapses of the hermaphrodite *C. elegans* connectome [23, 21, 20, 33] available from *WormWiring.org*[20]. All motifs tested occurred at least once in the *C. elegans* connectome. We discovered that the motifs encountered most frequently in one connectome tended to be the most frequent in other connectomes and their random-graph counterparts, but the absolute counts of each motif differed dramatically based upon the connectome and motif in question (**Figure 2**).

**Figure 2:**
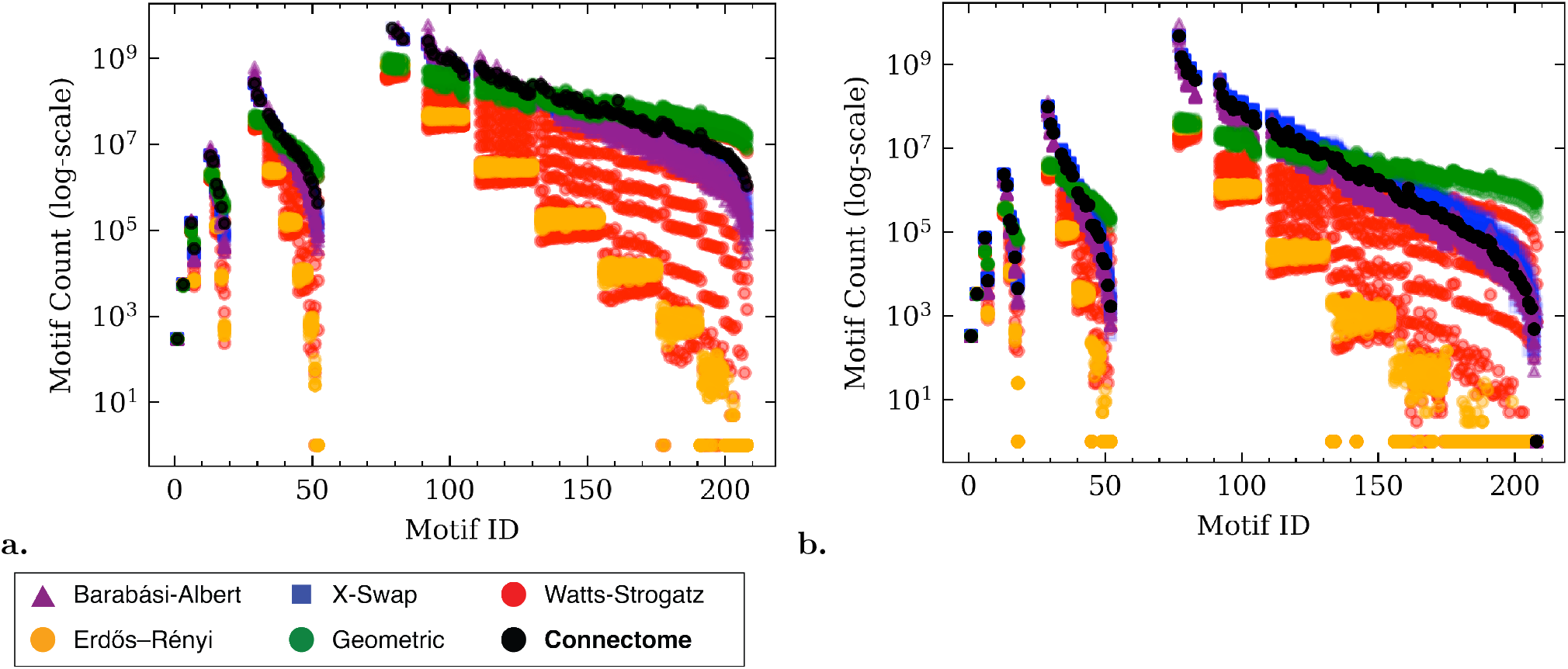
Undirected motif searches (monomorphisms) in connectomes and random graphs. **a & b:** The count of every undirected subgraph with six or fewer vertices (|*V*| ≤ 6). Each motif was counted in the connectomes, as well as in each of the random graph models calibrated to match the density of each connectome. The *x*-axis is the motif ID from the *Atlas of Graphs* text (e.g. all points on the line *x* = 7 represent the undirected triangle motif). Gaps along the *x*-axis indicate omitted motifs (i.e., those with multiple connected components). **a. Comparing***C. elegans* **with random graphs calibrated to***C. elegans* **density.** Due to its higher density, the *C. elegans* graph has higher motif counts than those of the MICrONS graph, despite a lower vertex count. X-swap motif counts (blue) closely match those of the original connectome (black). In contrast, Erdős–Rényi approximations (yellow) are very poor predictors of true connectome motif count, and always under-estimate motif counts in the original connectome. The parameter-space of the Watts-Strogatz model (red) leads to a wide range of motif count predictions, some as low as those of the Erdős–Rényi model. **b. Comparing MICrONS with random graphs calibrated to MICrONS density.** Like the *C. elegans* results in **a**, Erdős–Rényi approximations always underestimate the number of motifs, for all motif graphs we searched. The same motifs (*x* = 77, *x* = 78, etc) occur with the highest frequency in the MICrONS graph as in the *C. elegans* graph. Motif *x* = 208, the fully-connected graph on six nodes, appears many thousands of times in the *C. elegans* graph, but does not occur at all in the MICrONS connectome. Some models, like X-swap and Erdős–Rényi, likewise predict zero *K*_6_ motif occurrences. Others, like the geometric model (green), erroneously overpredict the number of expected *K*_6_ subgraphs. Full-resolution copies of this graphic are available online (see *Supplemental Materials*).

We also discovered that the distribution of undirected motifs (black datapoints, **Figure 2**) followed a similar curve trajectory for many brain graph datasets including the two shown here, though the parameters of that curve varied across connectomes. Queries from this set of experiments were run using Neo4j, NetworkX, and GrandIso executors, as convenient, leveraging our ability to seamlessly switch between executors when using DotMotif. Random graph parameters selected for these experiments are explained in greater depth in *Methods*. All data and results from this study are available as described in *Supplemental Materials*. These experiments may aid in the design or selection of random graph models to approximate a connectome graph, and in the design or selection of query graphs when studying reconstructed connectomes.

### 3.4 Directed three-node motif searches in MICrONS and *C. elegans*

Recent work has investigated the relative prevalence of directed three-node motifs in brain graphs and the importance of such motifs for local computation [10, 34, 35, 36]. Here we count all unique, directed, three-node motifs in the *MICrONS Phase 1* and *C. elegans* connectomes. Lower-density three-node motifs appeared with greater frequency (due to the larger number of embeddable permutations. Despite similar neuron counts, the connectomes differed dramatically in both absolute count and distribution shape, with many directed motifs occurring frequently in the *C. elegans* graph but few to no times in the MICrONS graph (**Figure 3**).

**Figure 3:**
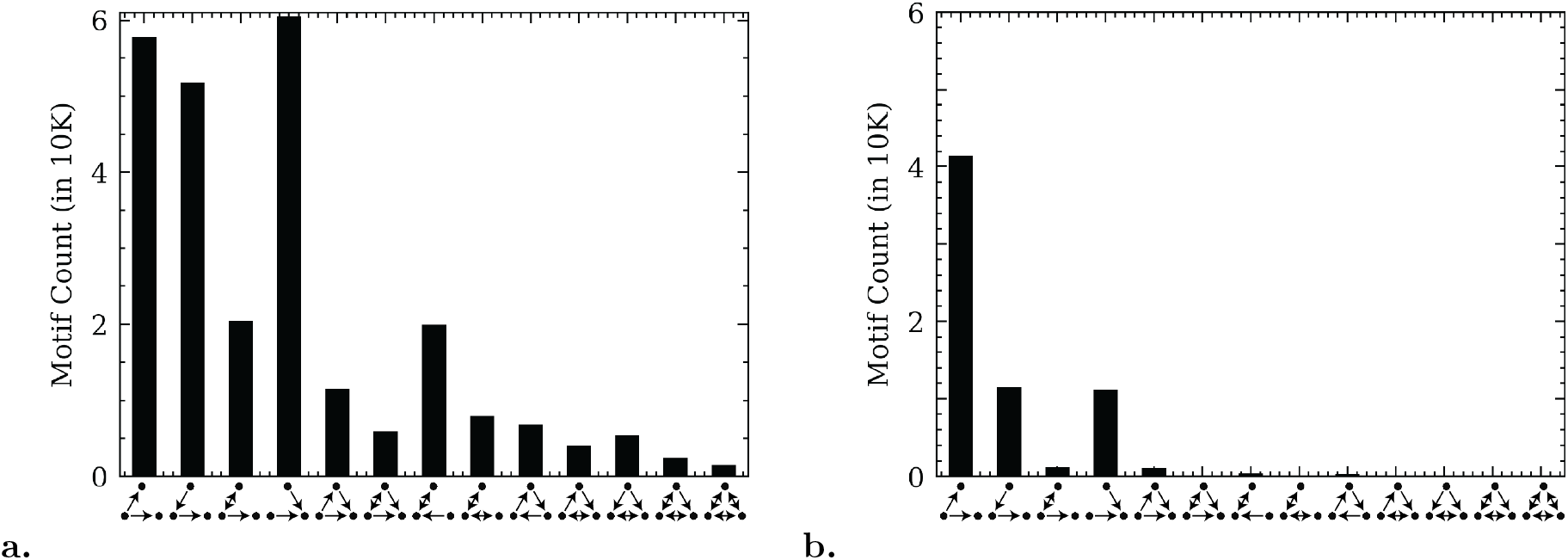
Quantities (monomorphisms) of directed three-node motifs from the C. elegans chemical synapse connectome and the MICrONS v185 graph. **a.***C. elegans* **directed three-node motif counts.** The most commonly encountered directed three-node motif is the “fan-in” motif, where two neurons converge to a single downstream target. All directed three-node motifs occur at least once. **b. MICrONS v185 directed three-node motif counts.** The most common motif is the “fan-out” motif, where one neuron synapses onto two downstream targets. Several three-node motifs appear infrequently or do not appear at all. This illustrates differences between the connectivity patterns of a complete connectome such as the worm’s and a mammalian brain region that is expected to be largely feed-forward.

These simple motifs serve as a useful frame for further investigations. Though there are only thirteen three-node directed graphs, exhaustively searching for larger directed motifs quickly becomes computationally infeasible (there are 199 connected four-node directed graphs, 9364 connected five-node directed graphs, and over 1.5 million connected six-node directed graphs). It is possible now to search specifically for larger supergraphs of *only* motifs which occur at least once in this study, greatly reducing the search space of future exhaustive motif searches in these connectomes. Unlike the distributions of undirected motifs, the relative distribution of these directed motifs does not appear to follow a consistent shape between connectomes. This suggests that even these small motifs may hold useful knowledge about the nature of connectome graphs, such as distinguishing between feed-forward-dominated or feedback-dominated tissue.

### 3.5 Benchmarks

Users may find that different DotMotif executors suit the needs of different research questions. In order to illustrate these differences, we compared the performance of three DotMotif executors available for use in our downloadable Python module. The *GrandIsoExecutor* and *NetworkXExecutor* are both pure-Python, and are therefore desirable for use in constrained compute environments such as shared servers. The *Neo4jExecutor* requires a Neo4j database, which runs as a standalone executable. **Figure 4** illustrates that the novel *GrandIso*-based executor developed under this effort always outperforms the NetworkX executor, regardless of host-graph size. When choosing between Python-based and graph-database executors, there is a tradeoff between data-ingest rate and search speed. The *NetworkXExecutor* and *GrandIsoExecutor*, which convert DotMotif DSL syntax into a series of Python commands, are preferable for running graph queries on a small host graph, or for running the same queries on many host graphs, due to low startup-time overhead. The *Neo4jExecutor*, which converts DotMotif DSL syntax into a Cypher query for execution with the Neo4j graph database, has significant data-ingest overhead for the first query on a graph, and so it is not ideal for running individual motif queries. However, users may prefer a Neo4j executor when running multiple queries on the same host graph, when many users are running motif queries simultaneously on the same host, or if motif searches are likely to be repeated. For queries on host-graphs of sufficiently large size, the *Neo4jExecutor* startup overhead becomes trivial compared to the total runtime of the query, and for larger-than-memory host-graphs, the *Neo4jExecutor* and *NeuPrintExecutors* may be the only feasible options. For most inmemory exploratory data analysis, we recommend the use of the *GrandIsoExecutor*. DotMotif users may trivially switch between Executors without modifying their queries, as the DotMotif engine stores queries in an intermediate format after parsing and validation steps (**Figure 5**).

**Figure 4:**
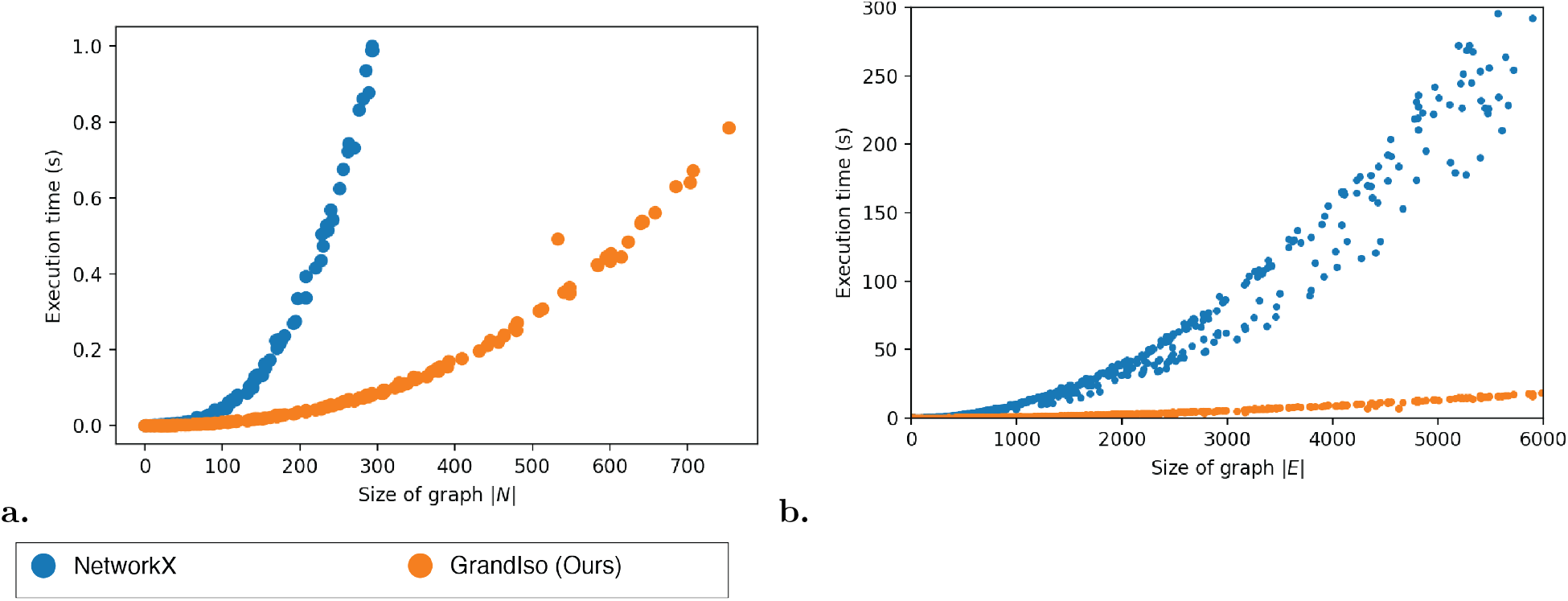
Comparison of wall-clock runtime when counting undirected motifs. All runtimes are measured during motif searches for undirected rectangles in Erdős-Rényi graphs of various sizes. All graph density parameters are 0.15. **a. Execution time as a function of node count.** Experiments were terminated if they exceeded one second of runtime. **b. Execution time as a function of edge count.** All experiments were performed on otherwise-idle hardware. Experiments were conducted on graphs with up to six thousand edges.

**Figure 5:**
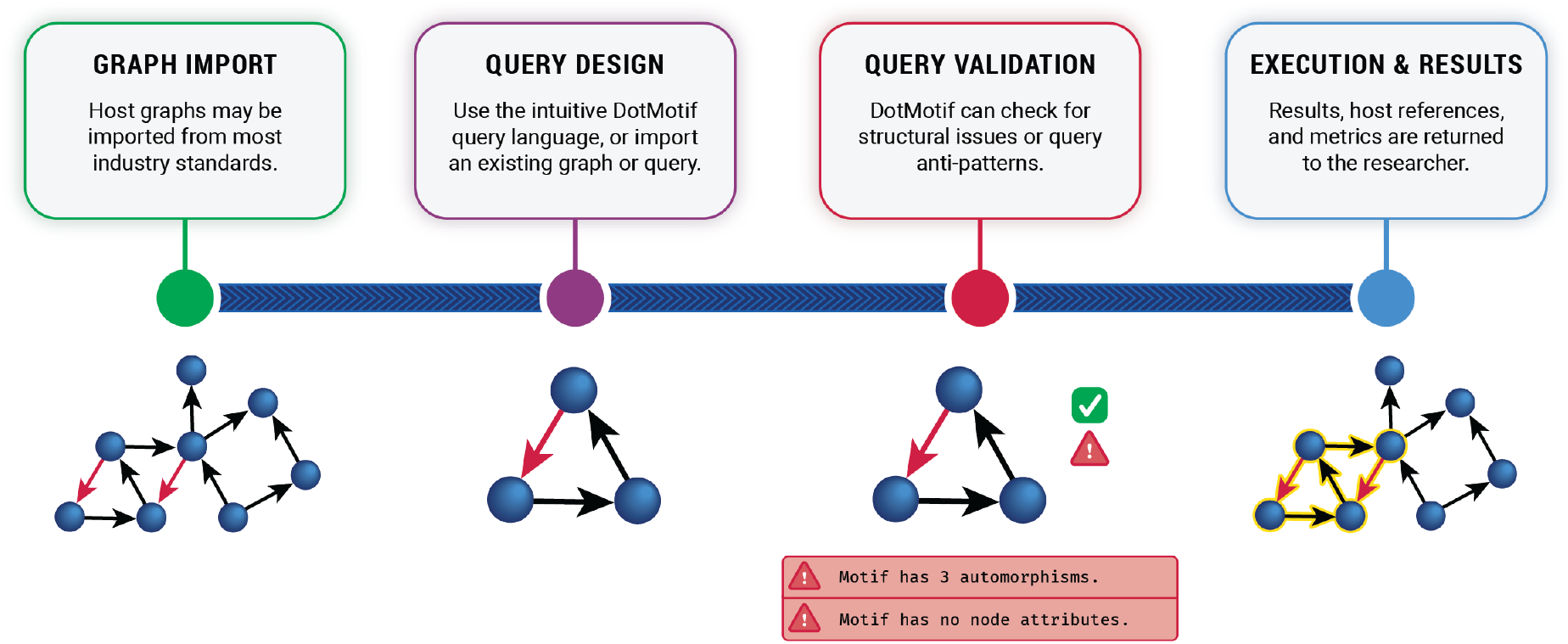
The DotMotif query execution process. **Graph Import**. A user may import a search graph (also commonly referred to as a *host* graph) from a variety of industry standard formats, including an edgelist CSV, GraphML, numpy adjacency matrix, and any format supported by the NetworkX library. Furthermore, DotMotif is compatible with graphs stored in neuPrint schemas in a Neo4j database [6]. **Query Design.** The user may select from a library of pre-built motifs, or write a novel query. The query may be written in the DotMotif DSL, encoded as a NetworkX graph, or written as text directly in the target graph database query language, such as Cypher. A user may choose to save this query as another standalone file on disk for future use. The .motif file-format stores both the motif itself and provenance metadata such as the date of creation, comments associated with the motif, and author information. **Query Validation.** DotMotif supports several pre-execution validation steps in order to fail quickly when asked to perform an impossible or self-contradictory query. Several other optional validators may be invoked to check a motif for biological feasibility and to warn the user if a potential error is detected. **Execution & Results.** The user may choose from the available DotMotif Executors in order to use the most appropriate subgraph matching tool for the compute resources available. No further query modification is required in order to run the same motif on several different Executors. A user may also choose to simply use DotMotif as a query executor without leveraging the query validation or query design tools.

## 4 Methods

In developing DotMotif, we aimed to ensure that our software was both accessible — intuitive for new users but powerful enough for power-users — as well as sufficiently computationally efficient to execute most common queries on commodity hardware. In the interest of research flexibility, we also required that the software adapt to best utilize available hardware resources. For this reason, DotMotif includes several *executors*, each of which leverage a different graph analysis technology. Executors are discussed in greater depth in the *Benchmarks* section.

### 4.1 Formal definition of the subgraph search task

Here we consider the ideas of *subgraph isomorphism* and *subgraph monomorphism*, which have slightly varying definitions in the graph theory literature. When we refer to a *subgraph*, we always refer to a *node-induced* subgraph, which is inline with the usage in popular subgraph isomorphism research, including research on the commonly used VF2 algorithm [37, 38, 39]. Given graphs *G* = (*V*_1_*, E*_1_) and *H* = (*V*_2_*, E*_2_), we say *G* is isomorphic to *H* if there exists a bijection *f* : *V*_1_ → *V*_2_ such that {*u, v*} ⊆ *E*_1_ if and only if *f* (*u*)*, f* (*v*) ⊆ *E*_2_ [40]. An extension of this idea is that a subgraph isomorphism exists between *G* and *H* if and only if there exists a subgraph *G^′^* of *G* such that *G^′^* is isomorphic to *H*. A monomorphism loosens the bijection requirement to simply injection, meaning that the match in the host graph may contain extra edges.

DotMotif finds all subgraphs in a search graph that match a query graph, where the term *match* used here means that one of the above described mappings exists. By default, DotMotif will perform a search for all subgraphs that are monomorphic to the query subgraph. A user may choose to search for exact matches only, in which case DotMotif identifies subgraph *isomorphisms*, rather than *monomorphisms*. This behavior is controlled by a user toggle during the construction of a query.

### 4.2 Optimizations to the subgraph isomorphism task in Python

For the purposes of quick iteration upon neuroscience hypotheses, our team required a high-speed subgraph isomorphism algorithm that was both pure Python — in order to reduce the barrier to entry for a fresh installation — as well as optimized to minimize CPU and memory usage. To satisfy this need, we developed a subgraph monomorphism search implementation that performs favorably against the standard NetworkX implementation (Fig 4). Our pure-Python implementation matches the results of the NetworkX *GraphMatcher* monomorphism detection module, with significantly lower CPU and memory overhead. Additionally, while many modern subgraph iso- and monomorphism implementations rely on a directed, acyclic graph state-space representation, our implementation stores its state space in a one-dimensional queue (**Algorithm 1**). In future work, we expect that due to this property, our algorithm may be trivially parallelized to operate on multiple cores at once.

### 4.3 Optimized interfaces for common graph libraries

Many existing research questions have been explored with RAM-sized graphs in mind, and have utilized popular Python graph libraries such as NetworkX or IGraph [17, 41], or emerging libraries such as Networkit [42]. In order to easily transition these algorithms and analyses to graphs of larger size, we developed an open-source graph library that enables a user to write code using familiar graph APIs (e.g. NetworkX’s nx.DiGraph), but which automatically converts these commands to run instead on any of a number of optimized, scalable backends, such as a graph database implemented in SQL or Amazon Web Services DynamoDB. This tool, alongside the subgraph iso- and monomorphism improvements detailed above, enabled us to achieve substantial performance speed-ups in performing the analyses in this paper. Links to documentation and the source code for this library are available in *Supplemental Materials*. All benchmarks listed here were performed on consumer laptop hardware with 16 gigabytes of RAM and a 3.1 GHz Intel Core i7 processor.

### 4.4 Comparison to Random Graph Models

One way to determine if a motif is significant within a real biological dataset is to compare its observed frequency to its expected frequency within a random graph model. There are many different random graph models, each with different properties and capturing a different aspect of real world data [43]. Here, we perform subgraph search on five different random graph models.

The Erdős-Rényi random model has parameters *n*, or the number of nodes, and *p*, or the probability that any of the potential 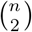 edges will independently exist within the graph [27]. The Erdős-Rényi model is easy to analyze and understand, and thus is included in this analysis. However, the edge independence assumption often fails for real datasets. Thus, other models are required to provide further context.

#### Algorithm 1: Our approach: Subgraph search

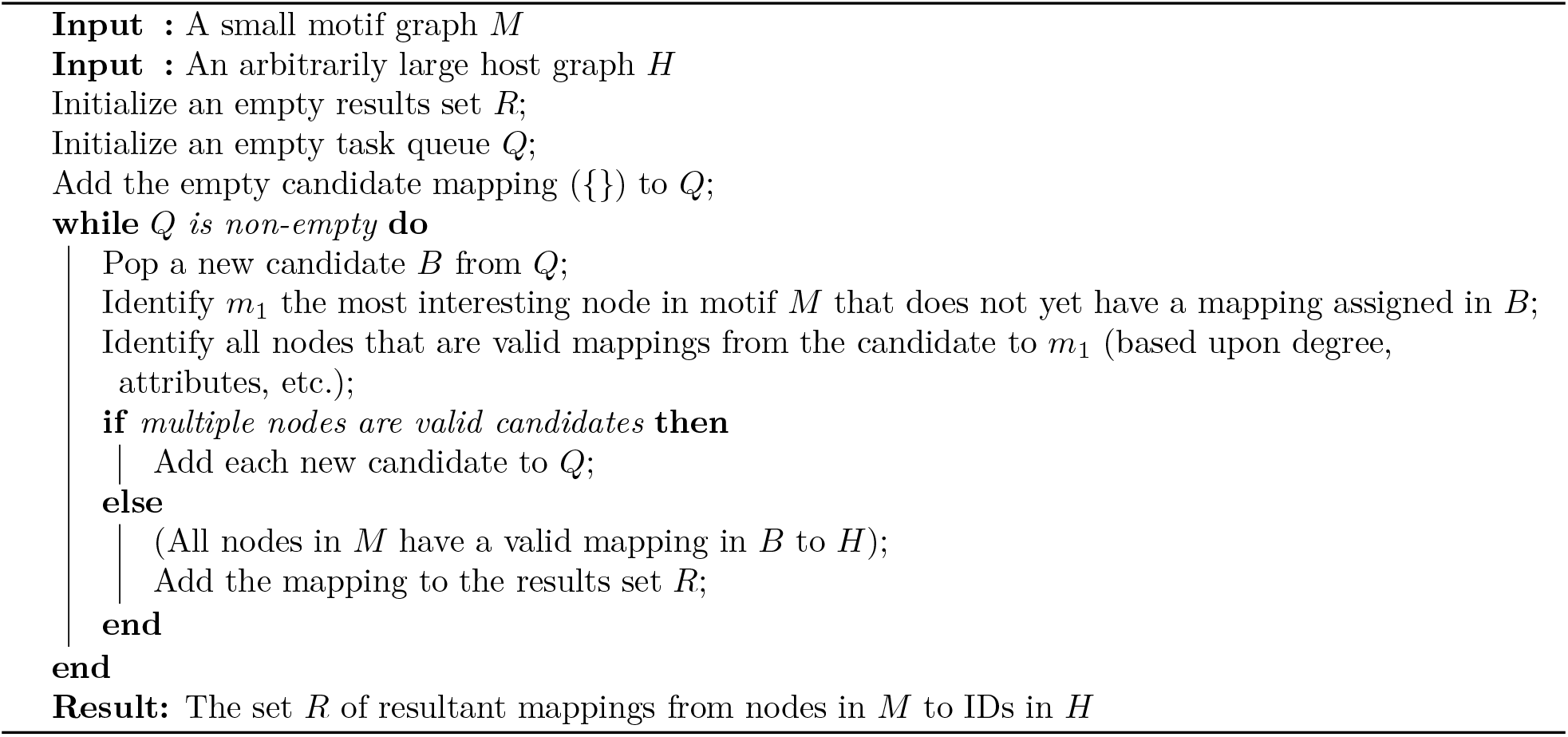

Geometric random graphs are desirable because they naturally capture spatial relationships that occur within the brain [44]. Nodes are uniformly distributed within a region and two nodes are adjacent if they fall within a radius *r*, which is an adjustable parameter [28]. This model, however, does not capture long-range neural connections, resulting in some spatial relationships that are not captured by the point cloud approach of the geometric model.

The Watts-Strogatz model addresses the edge dependence issue more directly, recognizing that many real world networks have high clustering coefficients, which is a measure of the extent to which two adjacent nodes have similar neighbors [29]. The Barabási-Albert model is a preferential-attachment model that ensures a power-law degree distribution, making it a scale free graph model [30, 43].

The degree-preserving edge-randomization graph model implemented here, based upon efforts such as those in Refs. [45, 32], accepts a graph as input-parameter and performs edge-swaps between two randomly-selected edges {*u*_1_*, v*_1_} and {*u*_2_*, v*_2_} so that the resulting edges are {*u*_1_*, v*_2_} and {*u*_2_*, v*_1_}. This model preserves the in- and out-degree distribution, as well as the in-degree sequence. Here, this model serves as a control in order to determine the role of degree distribution when determining the prevalence of a motif.

Each of the described random models has a set of parameters resulting in different graph properties. These parameters may be selected to generate graphs “similar” to the real dataset in question. Similarity can mean different things and we try to match the models to a characteristic that is closely tied to motif search. One obvious choice is to match the graph density 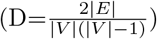 of each estimated connectome because density is a metric that can be more easily tweaked across models. Furthermore, a density that is too high or too low can result in inflated or underestimated subgraph counts, respectively [46]. Other metrics besides graph density may be considered, such as the degree distribution. Regardless of the metric, the parameters selected for a random graph model will differ for each connectome being analyzed. More specifically, differences in density for *C. elegans* and MICrONS datasets will result in differently parametrized random graph models. By performing subgraph search across these models tuned or calibrated to some characteristic of the real dataset, we narrow the problem space of what drives a given motif to occur at a given frequency. By understanding which models closely approximate certain phenomena such as motif count, we may be able to better identify biological mechanisms that induce these patterns of connectivity in the connectome.

### 4.5 Architecture

DotMotif is comprised of three submodules: A *parser* module, an *optimizer* module, and finally an *executor* module. In order of use, the *parser* module is responsible for converting the DotMotif domain-specific language into an in-memory representation. The *optimizer* module is then responsible for converting and simplifying the in-memory motif into its simplest possible representation. The *optimizer* module may also optionally check for violations of biological priors, in a validation step. Finally, the *executor* module converts the optimized motif into a query that can be submitted to a graph analysis tool. Each executor is responsible for generating its own target-specific queries. (For example, the *Neo4jExecutor* generates Cypher queries, and the *NetworkXExecutor* converts queries to a sequence of Python commands.) Through the coordination of these submodules, DotMotif provides a framework for posing and answering complex graph queries.

#### 4.5.1 DotMotif Domain-Specific Language Parser

We identified an impedance mismatch between the flexibility of common query languages (such as Cypher [7]) and the needs of the research community, where many research questions require overwhelmingly complex or verbose queries. We opted to develop a domain-specific query language to aid in the construction of queries. This enables research-driven query design agnostic to the underlying frameworks.

The DotMotif domain-specific language (DSL) borrows from DOT-like syntax [47] as well as from SQL-like syntax in order to expose a succinct and user-friendly query language. For example, the simple query A -> B will return a list of all edges in the complete graph (in other words, the list of all subgraphs of *G* = (*V, E*) that comprise one edge from a node *A* ∈ *V* to node *B* ∈ *V*). In order to make queries understandable and maintainable, the DotMotif DSL supports “macros,” or composable building-blocks that can be combined to generate more complicated queries (**Figure 6c**). These macros minimize “boilerplate” syntax without leading to duplicate notation (**Figure 6d**). Further illustrations of the DotMotif DSL syntax are available in **Figure 7a-b**.

**Figure 6:**
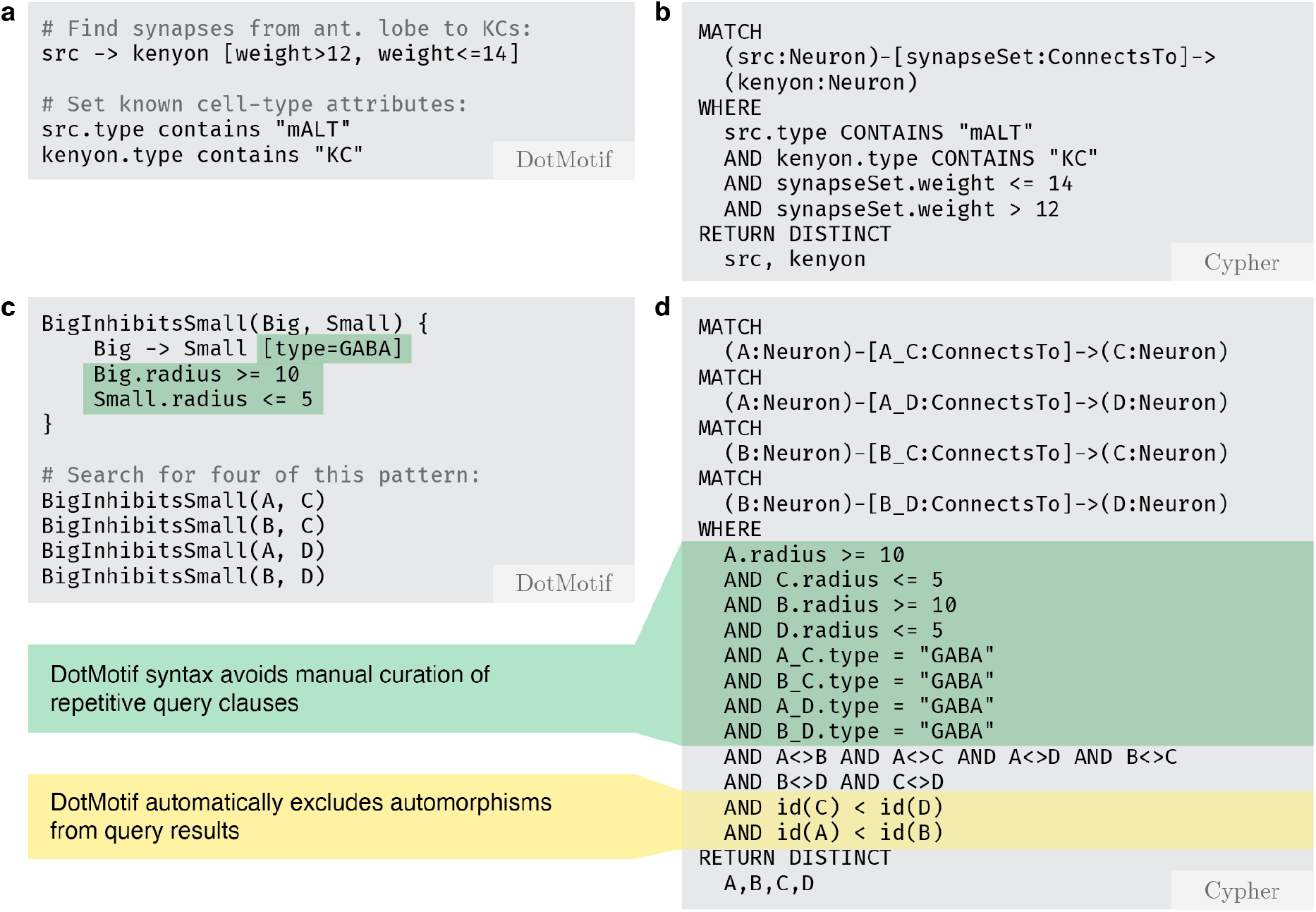
A comparison of DotMotif query syntax and the equivalent queries when transpiled to Cypher using the Neo4jExecutor. **a.** A simple query to find inputs to Kenyon cells (*KC)* from the medial antennal lobe tracts (*mALT*) in the *Hemibrain* dataset. Note that DotMotif comments are notated with the hash character. **b.** The equivalent query as panel ***a***, when converted to the Cypher query language for use with *neuPrint* systems. Even in a small query such as this, DotMotif syntax tends to be more succinct and readable. **c.** A simple DotMotif query for a repeated pattern with node and edge constraints. The motif includes a *macro* called BigInhibitsSmall, which establishes a connection between two neurons with a *type* edge-attribute of “GABA”, and with *radius* nodes attribute constraints on both neuron participants. This macro is reused multiple times in the final motif construction in order to avoid repetitive code. DotMotif automatically infers that nodes *A* and *B* are isomorphic (i.e. interchangeable), and that *C* and *D* are isomorphic. **d.** DotMotif converts the query in panel ***c*** to the Cypher code in **d**. The equivalent Cypher query is substantially longer and harder to maintain or edit, even in the case of this quite simple motif. It also requires explicit notation to avoid reporting duplicate motif automorphisms.

**Figure 7:**
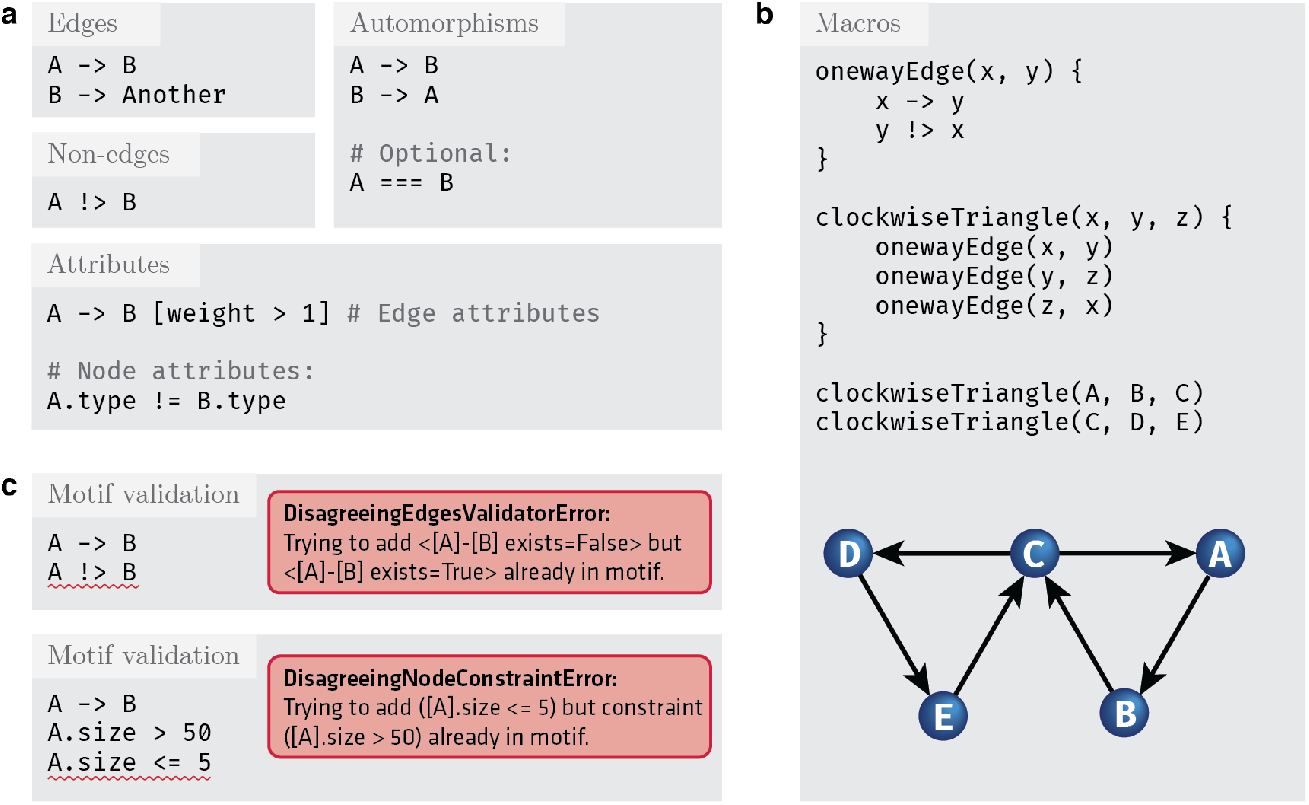
Examples of the DotMotif query language. **a.** Features of the DotMotif domain-specific language syntax. Nodes are connected by directed edges, notated with an arrow operator. A user can specify whether a certain edge must *not* exist by using the non-edge operator. Both edges as well as nodes may be assigned constraints or attributes. Edge attributes are nested in square brackets on the same line as the edge notation, and node attributes are notated with ‘dot’ notation on their own line. A user can explicitly indicate that two nodes are interchangeable, and DotMotif will only return one representative of that automorphism group. **b.** Example use of nested macros to construct a complex motif from simple building blocks. In this motif example, *x*, *y*, and *z* serve as macro arguments. These variables, similar to local function arguments in other programming languages, are only used within the macro, and are not participants in the motif. Nodes with names *A* – *E* serve as participants in the motif. Note that macros may be nested by calling one from within the body of another. **c.** Examples of motifs that fail validation. The DotMotif query validation step reduces the likelihood of spending computational resources on impossible or biologically unfeasible queries. In the first example, validation fails because an edge has already been added to the motif but a new line (red underline) conflicts with this requirement. In the second example, validation fails because the condition is impossible (no value for `A.size` can be both greater than 50 as well as less than or equal to 5). These validation failures serve as early warning-signs for a researcher to see that a query will fail if executed.

By default, DotMotif returns a single representative element of each automorphism group in the result set. An *automorphism* is an isomorphism from a graph onto itself [40]. Returning a single representative of the automorphism group avoids over-counting motifs in the search graph due to motif symmetries. The ability to count only one representative of this group is useful to many common neuroscience questions, but it is not easily accomplished with conventional graph tools. A DotMotif user may also explicitly specify that two nodes in a motif are interchangeable with the automorphism operator (**Figure 7a**). Whether the user uses automatic query automorphism detection or notates it manually, the DotMotif optimizer will enrich the query motif to lower the space of possible matches, and thus return a result more rapidly. DotMotif considers both semantic as well as syntactic feasibility when automatically determining automorphism mappings [38].

This language is formally defined in Extended Backus–Naur Form [48, 49] by the grammar file referenced in *Supplemental Materials*. This syntax is visualized using the railroad-diagram convention in **Figure 8**.

**Figure 8:**
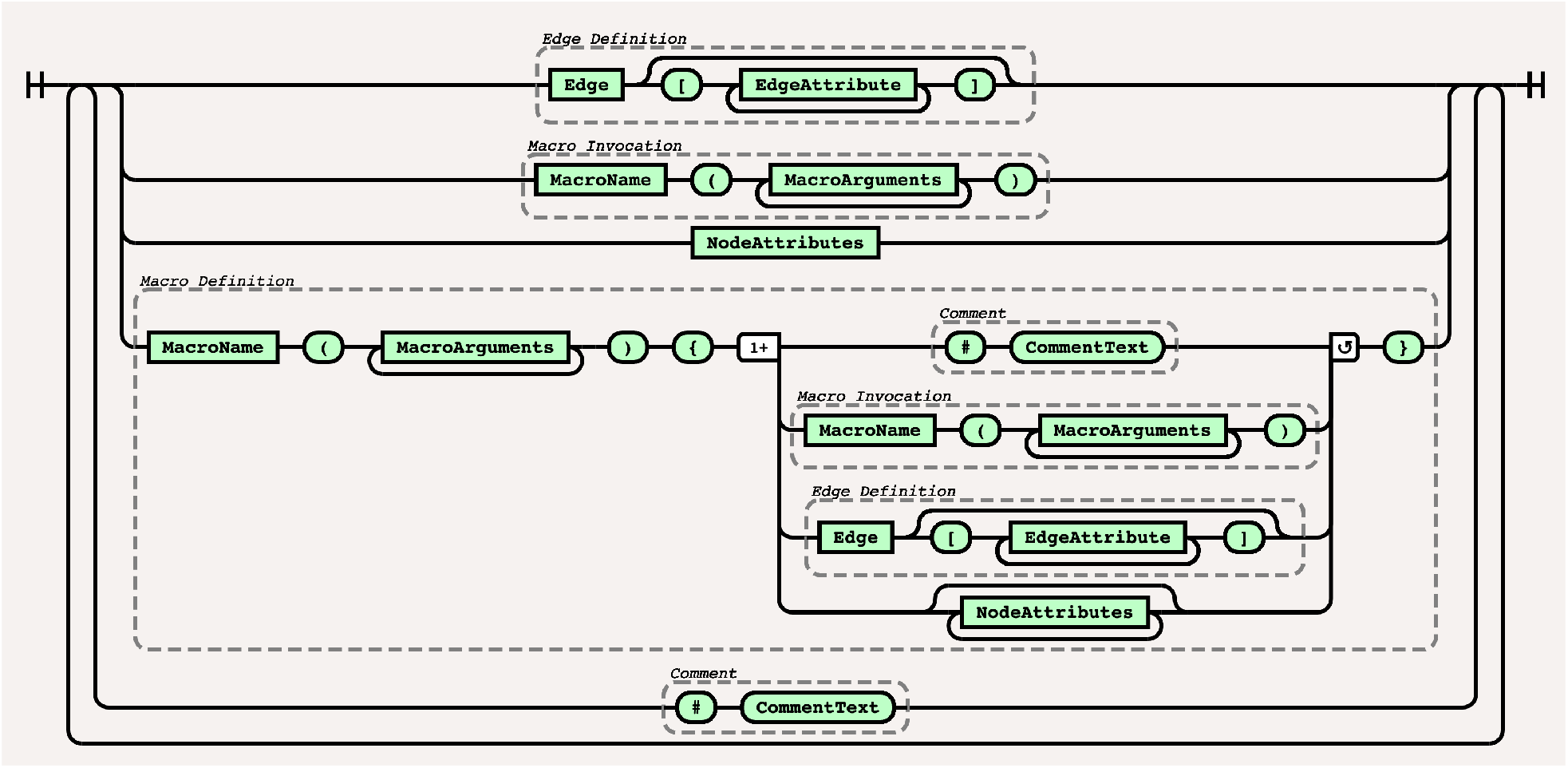
The DotMotif Syntax in railroad-diagram form. The DotMotif DSL is a whitespace-agnostic language with hash-symbol comments and curly-brace bracketed macros. Edges are defined with ->-arrowlike syntax; edge attributes may be listed in JSON-like syntax. Node attributes may be listed in object dotnotation syntax. For example usage of this syntax system, refer to *Architecture*. This syntax is formally defined in Extended Backus-Naur Form. Both this formal definition as well as the language specification will be made available as per *Supplemental Materials*. This figure was generated with the *railroad-diagrams* Node.js package [50].

#### 4.5.2 Query optimization and validation

After a query has been parsed and ingested, it is passed to an optimization stage in which the query is reduced to a simplified form in order to improve the speed of query execution. Optionally, a user may also enable the automatic detection of impossible structures — or even biologically implausible structures — in a *Validation* step (**Figure 7c**).

These verbose error messages are intended to warn a user quickly and clearly if a graph query is unlikely or impossible, rather than allowing the user to proceed with a long-running query that is destined to fail. In contrast, the equivalent Cypher command running in Neo4j will still go through the process of execution on the full host graph, but will return no results. DotMotif includes the option to add validators for *biological plausibility*. These options allow the user to prohibit motifs that — for example — require inhibition and excitation to be performed by the same neuron. These validators, like all validators in the DotMotif package, are both optional as well as modifiable.

#### 4.5.3 Query execution

A parsed, optimized, and validated query can now be evaluated against a large graph. Where necessary, it is possible to run these queries fully in-memory on consumer hardware. DotMotif includes both pure Python as well as graph-database query executors. There is no difference between the results returned from each executor, and so they may be used interchangeably, depending upon the needs and parameters of the experiments and environments.

In order to take advantage of graph database technology, we have implemented a *Neo4jExecutor*, which leverages the Neo4j [7] database and its built-in subgraph match detection algorithm, an implementation of the *VF2* algorithm [37, 38]. In order to run queries in this environment, our software converts the optimized query to *Cypher*, the database query language used natively by Neo4j. If a user does not have a running Neo4j database, our library also includes routines to provision a Docker container [51] and quickly ingest the data in an appropriate format for Neo4j to use. A user may also direct queries to a running neuPrint [6] database. A list of all available executors is available in **Table 2**.

**Table 2:**
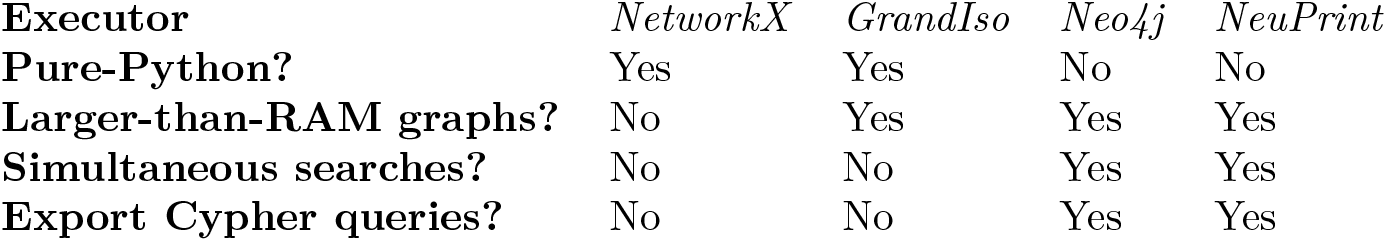
*A comparison of DotMotif executors*. All executors are capable of running any of the queries in this paper. Pure Python executors are desirable due to their low configuration and installation overhead, though they tend to be slower than graph databases. Searches in larger-than-RAM graphs are partially supported in the *GrandIso* executor with the use of the *Grand* library discussed in *Optimized interfaces for common graph libraries*. Graph databases support this by default. Both *neuPrint* and *Neo4j* run as a separate service, and both accept simultaneous client connections, whereas the Python executors cannot. The user can use these executors interchangeably, and may find that different executors work better for specific types of motif searches.

## 5 Discussion

Though modern connectomics research commonly interprets neural connectivity as a network, the field currently suffers from a lack of accessible, performant graph analysis and subgraph-search tools. In this work, we present DotMotif, a combined domain-specific language and interface to powerful graph-search tools. Our hope is that this tool and others like it will reduce the barrier-to-entry for researchers unacquainted with graph theory or graph databases, and will enable researchers to interrogate increasingly common connectome datasets with ease. Though the undirected graphs from the graph atlas study may not directly provide answers to open questions in neuroscience, this broad search can narrow down where it may be most impactful to dig deeper, perhaps leading to more interesting directed graph searches or neural simulations over smaller directed versions. Future work may include random assignment of edge-direction in order to perform an unbiased search across the space of all possible directed graphs, akin to other random-graph searches [8]. We intend to further explore other metrics beyond density for random graph model calibration in the future.

Through these directed and undirected motif studies, it may also be possible to build a heuristic to predict motif-counts in a connectome using a small number of simpler (i.e. lower computational cost) motif searches, and avoid explicit counting altogether. For example, if the goal of a research study were to determine if the actual count of a motif in question aligns with predictions for a connectome of a certain modality, size, species, and brain region, such a model would circumvent the need to search in random graph models, and would instead enable a simpler, analytic approach.

Recent neuroscience research [52, 34, 53, 10, 54] has provided many valuable explanations for neuron-to-neuron connectivity rules. We intend for these tools to offer a quantitative way to expand these rules to larger numbers of neurons, and to free the neuroscience community to explore new questions in the connectomics domain. Similarly, several efforts have begun to map subneuron connectivity patterns, identifying recurring motifs in the relationships of multiple synapses between the same neurons [55, 56, 57]. We believe that the systems presented here will provide a substrate upon which to build not only nanoscale connectivity research but also subcellular motif work and meso- to macro-scale connectomics, through modalities such as X-ray microtomography or MRI [58, 16, 59].

Despite many of the simplifications and optimizations to the subgraph monomorphism task mentioned here, this task in general is NP-complete [60, 39, 61, 62]. Even in a relatively small connectome such as that of *C. elegans*, certain sufficiently common or poorly-optimized motif searches may still remain infeasible to run on consumer hardware. In our experience, the addition of further constraints, such as specifying edge direction, specifying node- and edge-attributes, or searching for query graphs with nodes of high degree, may reduce the execution time of complex queries; but other seemingly helpful modifications, such as increasing the diameter of a query graph, may counter-intuitively increase runtime quite substantially. As a result of such longer query runtimes or more compute-hungry queries, researchers will be able to run fewer queries per study.

In our ongoing work, we hope to reduce this barrier by developing more rigorous automated query optimizations and by publishing connectome query results for reuse. We will introduce and refine new validators that aid DotMotif query designers in identifying qualities of a query graph that may lead to long runtimes, and we hope that the atlas results published above may shed light on how to select query graphs for further study in a dataset. We intend to disseminate an open-data *motif encyclopedia* so that long-running or complex queries may be run on a dataset once and then shared with the community.

We also note that faster implementations of the motif-search problem — relevant to connectomics, pharmaceutical research, and other domains — may pave the way to faster and more efficient scientific discovery.

The field of connectomics is at an inflection point as datasets continue to grow in size, as technology and neuroscience provide avenues to create and study large datasets at unprecedented scales beyond the analysis capabilities of a single lab. Open, free, and public datasets, as well as accessible and affordable tools to understand those public data, are of paramount importance.

We are releasing the DotMotif codebase, as well as all demonstration code and data in this manuscript, as open-source and open-data tools to support community discovery. We invite collaborators to share questions that allow us extend DotMotif to test scientific hypotheses.

## 6 Supplemental Materials

Code for the DotMotif Python library, the example snippets in this paper, and all data and code for the experiments run in *Results* are available online at *bossdb.org/tools/dotmotif*.

Code and documentation for the improved subgraph isomorphism library is available online at *bossdb.org/tools/grandiso*, and the graph-library abstraction tool discussed in *Optimized interfaces for common graph libraries* is available at *bossdb.org/tools/grand*.

## 7 Acknowledgements

The authors would like to thank Konrad Kording for advice and review of this manuscript. We would like to thank the teams that generated the connectome data used in this analysis.

Research reported in this publication was supported by the National Institute of Mental Health of the National Institutes of Health under Award Numbers R24MH114799 and R24MH114785. The content is solely the responsibility of the authors and does not necessarily represent the official views of the National Institutes of Health. This work was also completed with the support of JHU/APL Internal Research Funding.

## Author contributions statement

J.M. and J.S. conducted the experiments and designed the software. E.R. and E.J. analyzed the results. B.W., D.B., and W.G.R. conceived and designed the experiments, analyzed the results, and secured funding for this work. All authors wrote and reviewed the manuscript.

## References

[1] D. S. Bassett, P. Zurn, and J. I. Gold, “On the nature and use of models in network neuroscience,” Nature Reviews Neuroscience, vol. 19, no. 9, p. 566–578, Jul 2018. [Online]. Available: http://dx.doi.org/10.1038/s41583-018-0038-8

[2] C. S. Xu, M. Januszewski, Z. Lu, S.-y. Takemura, K. J. Hayworth, G. Huang, K. Shinomiya, J. Maitin-Shepard, D. Ackerman, S. Berg, and et al., “A Connectome of the Adult Drosophila Central Brain,” biorXiv, Jan 2020. [Online]. Available: http://dx.doi.org/10.1101/2020.01.21.911859

[3] R. Burns, E. Perlman, A. Baden, W. G. Roncal, B. Falk, V. Chandrashekhar, F. Collman, S. Seshamani, J. Patsolic, and K. Lillaney, “A Community-Developed Open-Source Computational Ecosystem for Big Neuro Data,” arXiv preprint arXiv:1804.02835, 2018.

[4] T. Hočevar and J. Demšar, “Combinatorial algorithm for counting small induced graphs and orbits,” PLOS ONE, vol. 12, no. 2, p. e0171428, Feb. 2017. [Online]. Available: http://dx.doi.org/10.1371/journal.pone.0171428

[5] L. K. Scheffer, “Graph Properties of the Adult Drosophila Central Brain,” biorXiv, May 2020. [Online]. Available: http://dx.doi.org/10.1101/2020.05.18.102061

[6] J. Clements, T. Dolafi, L. Umayam, N. L. Neubarth, S. Berg, L. K. Scheffer, and S. M. Plaza, “neuPrint: Analysis Tools for EM Connectomics,” biorXiv, Jan. 2020. [Online]. Available: http://dx.doi.org/10.1101/2020.01.16.909465

[7] D. Fernandes and J. Bernardino, “Graph Databases Comparison: AllegroGraph, ArangoDB, InfiniteGraph, Neo4J, and OrientDB.” in DATA, 2018, pp. 373–380.

[8] O. Sporns and R. Kötter, “Motifs in brain networks,” PLOS BIOL, p. 369, 2004.

[9] V. B. Mountcastle, “Modality and Topographic Properties of Single Neurons of Cat’s Somatic Sensory Cortex,” Journal of Neurophysiology, vol. 20, no. 4, p. 408–434, Jul. 1957. [Online]. Available: http://dx.doi.org/10.1152/jn.1957.20.4.408

[10] N. L. Turner, T. Macrina, J. A. Bae, R. Yang, A. M. Wilson, C. Schneider-Mizell, K. Lee, R. Lu, J. Wu, A. L. Bodor, and et al., “Multiscale and multimodal reconstruction of cortical structure and function,” biorXiv, Oct. 2020. [Online]. Available: http://dx.doi.org/10.1101/2020.10.14.338681

[11] N. Pospelov, S. Nechaev, K. Anokhin, O. Valba, V. Avetisov, and A. Gorsky, “Spectral peculiarity and criticality of a human connectome,” Physics of Life Reviews, vol. 31, p. 240–256, Dec. 2019. [Online]. Available: http://dx.doi.org/10.1016/j.plrev.2019.07.003

[12] E. P. Reilly, J. S. Garretson, W. R. Gray Roncal, D. M. Kleissas, B. A. Wester, M. A. Chevillet, and M. J. Roos, “Neural reconstruction integrity: A metric for assessing the connectivity accuracy of reconstructed neural networks,” Frontiers in neuroinformatics, vol. 12, p. 74, 2018.

[13] E. P. Reilly, E. C. Johnson, M. J. Hughes, D. Ramsden, L. Park, B. Wester, and W. Gray-Roncal, “Connecting Neural Reconstruction Integrity (NRI) to Graph Metrics and Biological Priors,” in Complex Networks XI. Springer, 2020, pp. 182–193.

[14] O. Sporns, “Structure and function of complex brain networks,” Dialogues in Clinical Neuroscience, vol. 15, pp. 247–262, 2013.

[15] L. W. Swanson and J. W. Lichtman, “From Cajal to Connectome and Beyond,” Annual Review of Neuroscience, vol. 39, no. 1, p. 197–216, Jul. 2016. [Online]. Available: http://dx.doi.org/10.1146/annurev-neuro-071714-033954

[16] D. S. Bassett and O. Sporns, “Network neuroscience,” Nature Neuroscience, vol. 20, no. 3, p. 353–364, Feb. 2017. [Online]. Available: http://dx.doi.org/10.1038/nn.4502

[17] A. A. Hagberg, D. A. Schult, and P. J. Swart, “Exploring Network Structure, Dynamics, and Function using NetworkX,” in Proceedings of the 7th Python in Science Conference, G. Varoquaux, T. Vaught, and J. Millman, Eds., Pasadena, CA USA, 2008, pp. 11–15.

[18] Google, “Cayley: An open-source graph database,” Jul. 2020. [Online]. Available: https://github.com/cayleygraph/cayley

[19] R. Wang, Z. Yang, W. Zhang, and X. Lin, “An Empirical Study on Recent Graph Database Systems,” Lecture Notes in Computer Science, p. 328–340, 2020. [Online]. Available: http://dx.doi.org/10.1007/978-3-030-55130-8_29

[20] S. J. Cook, T. A. Jarrell, C. A. Brittin, Y. Wang, A. E. Bloniarz, M. A. Yakovlev, K. C. Q. Nguyen, L. T.-H. Tang, E. A. Bayer, J. S. Duerr, and et al., “Whole-animal connectomes of both Caenorhabditis elegans sexes,” Nature, vol. 571, no. 7763, p. 63–71, Jul. 2019. [Online]. Available: http://dx.doi.org/10.1038/s41586-019-1352-7

[21] J. White, E. Southgate, J. N. Thomson, and S. Brenner, “The structure of the nervous system of the nematode C. elegans,” Philosophical transactions Royal Society London, vol. 314, pp. 1–340, 1986.

[22] L. Becker, A. L. Bodor, A. A. Bleckert, D. Brittain, J. Buchanan, D. J. Bumbarger, M. Castro, F. Collman, S. Dorkenwald, E. Froudarakis, D. Ih, N. Kemnitz, C. S. Jordan, K. Lee, Y. Li, R. Lu, N. M. da Costa, T. Macrina, G. Mahalingam, S. Popovych, R. C. Reid, J. Reimer, H. S. Seung, C. Schneider-Mizell, W. Silversmith, S. Suckow, M. Takeno, N. L. Turner, I. Tartavull, A. S. Tolias, R. Torres, A. M. Wilson, W. Wong, J. Wu, S.-C. Yu, A. Zlateski, and J. Zung, “MICrONS Layer 2/3 Data Tables,” Mar. 2020. [Online]. Available: https://doi.org/10.5281/zenodo.3710459

[23] T. A. Jarrell, Y. Wang, A. E. Bloniarz, C. A. Brittin, M. Xu, J. N. Thomson, D. G. Albertson, D. H. Hall, and S. W. Emmons, “The Connectome of a Decision-Making Neural Network,” Science, vol. 337, no. 6093, p. 437–444, Jul. 2012. [Online]. Available: http://dx.doi.org/10.1126/science.1221762

[24] S. Dorkenwald, N. L. Turner, T. Macrina, K. Lee, R. Lu, J. Wu, A. L. Bodor, A. A. Bleckert, D. Brittain, N. Kemnitz, and et al., “Binary and analog variation of synapses between cortical pyramidal neurons,” biorXiv, Dec. 2019. [Online]. Available: http://dx.doi.org/10.1101/2019.12.29.890319

[25] C. M. Schneider-Mizell, A. L. Bodor, F. Collman, D. Brittain, A. A. Bleckert, S. Dorkenwald, N. L. Turner, T. Macrina, K. Lee, R. Lu, and et al., “Chandelier cell anatomy and function reveal a variably distributed but common signal,” biorXiv, Apr. 2020. [Online]. Available: http://dx.doi.org/10.1101/2020.03.31.018952

[26] R. C. Read and R. J. Wilson, An Atlas of Graphs. USA: Oxford University Press, Inc., 2005.

[27] P. Erdös and A. Rényi, “On Random Graphs I,” Publicationes Mathematicae Debrecen, vol. 6, p. 290, 1959.

[28] D. Penrose, M. Penrose, and O. U. Press, Random Geometric Graphs, ser. Oxford studies in probability. Oxford University Press, 2003. [Online]. Available: https://books.google.com/books?id=RHvnCwAAQBAJ

[29] D. J. Watts and S. H. Strogatz, “Collective dynamics of “small-world” networks,” Nature, vol. 393, no. 6684, p. 440–442, Jun. 1998. [Online]. Available: http://dx.doi.org/10.1038/30918

[30] R. Albert and A.-L. Barabási, “Statistical mechanics of complex networks,” Reviews of Modern Physics, vol. 74, no. 1, p. 47–97, Jan. 2002. [Online]. Available: http://dx.doi.org/10.1103/RevModPhys.74.47

[31] M. E. J. Newman, “The Structure and Function of Complex Networks,” SIAM Review, vol. 45, no. 2, p. 200–202, Jan 2003. [Online]. Available: http://dx.doi.org/10.1137/S003614450342480

[32] E. S. Roberts and A. C. C. Coolen, “Unbiased degree-preserving randomisation of directed binary networks,” 2011.

[33] C. A. Brittin, S. J. Cook, D. H. Hall, S. W. Emmons, and N. Cohen, “Volumetric reconstruction of main Caenorhabditis elegans neuropil at two different time points,” biorXiv, Dec. 2018. [Online]. Available: http://dx.doi.org/10.1101/485771

[34] E. Gal, R. Perin, H. Markram, M. London, and I. Segev, “Neuron Geometry Underlies Universal Network Features in Cortical Microcircuits,” biorXiv, May 2019. [Online]. Available: http://dx.doi.org/10.1101/656058

[35] C. Curto, C. Langdon, and K. Morrison, “Robust motifs of threshold-linear networks,” 2019.

[36] A. J. Whalen, S. N. Brennan, T. D. Sauer, and S. J. Schiff, “Observability and Controllability of Nonlinear Networks: The Role of Symmetry,” Phys. Rev. X, vol. 5, p. 011005, Jan 2015. [Online]. Available: https://link.aps.org/doi/10.1103/PhysRevX.5.011005

[37] L. P. Cordella, P. Foggia, C. Sansone, and M. Vento, “An improved algorithm for matching large graphs,” in In: 3rd IAPR-TC15 Workshop on Graph-based Representations in Pattern Recognition, Cuen, 2001, pp. 149–159.

[38] L. Cordella, P. Foggia, C. Sansone, and M. Vento, “A (Sub)Graph Isomorphism Algorithm for Matching Large Graphs,” Pattern Analysis and Machine Intelligence, IEEE Transactions on, vol. 26, pp. 1367–1372, 11 2004.

[39] J. R. Ullmann, “An Algorithm for Subgraph Isomorphism,” Journal of the ACM (JACM), vol. 23, no. 1, p. 31–42, Jan. 1976. [Online]. Available: http://dx.doi.org/10.1145/321921.321925

[40] D. B. West et al., Introduction to graph theory. Prentice hall Upper Saddle River, 2001, vol. 2.

[41] G. Csardi and T. Nepusz, “The igraph software package for complex network research,” InterJournal, vol. Complex Systems, p. 1695, 2006. [Online]. Available: http://igraph.org

[42] C. L. Staudt, A. Sazonovs, and H. Meyerhenke, “NetworKit: A Tool Suite for Large-scale Complex Network Analysis,” 2015.

[43] J. Chung, E. Bridgeford, J. Arroyo, B. D. Pedigo, A. Saad-Eldin, V. Gopalakrishnan, L. Xiang, C. E. Priebe, and J. T. Vogelstein, “Statistical Connectomics,” Center for Open Science, Aug. 2020. [Online]. Available: http://dx.doi.org/10.31219/osf.io/ek4n3

[44] J. Stiso and D. S. Bassett, “Spatial Embedding Imposes Constraints on Neuronal Network Architectures,” Trends in Cognitive Sciences, vol. 22, no. 12, p. 1127–1142, Dec 2018. [Online]. Available: http://dx.doi.org/10.1016/j.tics.2018.09.007

[45] S. Maslov, “Specificity and Stability in Topology of Protein Networks,” Science, vol. 296, no. 5569, p. 910–913, May 2002. [Online]. Available: http://dx.doi.org/10.1126/science.1065103

[46] B. C. M. van Wijk, C. J. Stam, and A. Daffertshofer, “Comparing Brain Networks of Different Size and Connectivity Density Using Graph Theory,” PLoS ONE, vol. 5, no. 10, p. e13701, Oct. 2010. [Online]. Available: http://dx.doi.org/10.1371/journal.pone.0013701

[47] E. R. Gansner, E. Koutsofios, S. C. North, and K. phong Vo, “A Technique for Drawing Directed Graphs,” IEEE Transactions on Software Engineering, vol. 19, no. 3, pp. 214–230, 1993.

[48] J. W. Backus, “The syntax and semantics of the proposed international algebraic language of the Zurich ACM-GAMM Conference.” in IFIP Congress. Butterworths, London, 1959, pp. 125–131. [Online]. Available: http://dblp.uni-trier.de/db/conf/ifip/ifip1959.html#Backus59

[49] D. E. Knuth, “Backus Normal Form vs. Backus Naur Form,” Communications of the ACM, vol. 7, no. 12, p. 735–736, Dec. 1964. [Online]. Available: http://dx.doi.org/10.1145/355588.365140

[50] T. Atkins-Bittner, “Railroad-diagram Generators,” https://github.com/tabatkins/railroad-diagrams, 2020.

[51] D. Merkel, “Docker: lightweight linux containers for consistent development and deployment,” Linux journal, vol. 2014, no. 239, p. 2, 2014.

[52] R. F. Betzel, A. Avena-Koenigsberger, J. Goñi, Y. He, M. A. de Reus, A. Griffa, P. E. Vértes, B. Mišic, J.-P. Thiran, P. Hagmann, and et al., “Generative models of the human connectome,” NeuroImage, vol. 124, p. 1054–1064, Jan. 2016. [Online]. Available: http://dx.doi.org/10.1016/j.neuroimage.2015.09.041

[53] M. Schröter, O. Paulsen, and E. T. Bullmore, “Micro-connectomics: probing the organization of neuronal networks at the cellular scale,” Nature Reviews Neuroscience, vol. 18, no. 3, p. 131–146, Feb. 2017. [Online]. Available: http://dx.doi.org/10.1038/nrn.2016.182

[54] R. Milo, “Network Motifs: Simple Building Blocks of Complex Networks,” Science, vol. 298, no. 5594, p. 824–827, Oct. 2002. [Online]. Available: http://dx.doi.org/10.1126/science.298.5594.824

[55] J. L. Morgan, D. R. Berger, A. W. Wetzel, and J. W. Lichtman, “The Fuzzy Logic of Network Connectivity in Mouse Visual Thalamus,” Cell, vol. 165, no. 1, p. 192–206, Mar. 2016. [Online]. Available: http://dx.doi.org/10.1016/j.cell.2016.02.033

[56] J. L. Morgan and J. W. Lichtman, “An Individual Interneuron Participates in Many Kinds of Inhibition and Innervates Much of the Mouse Visual Thalamus,” Neuron, vol. 106, no. 3, p. 468–481.e2, May 2020. [Online]. Available: http://dx.doi.org/10.1016/j.neuron.2020.02.001

[57] A. M. Wilson, R. Schalek, A. Suissa-Peleg, T. R. Jones, S. Knowles-Barley, H. Pfister, and J. W. Lichtman, “Developmental Rewiring between Cerebellar Climbing Fibers and Purkinje Cells Begins with Positive Feedback Synapse Addition,” Cell Reports, vol. 29, no. 9, p. 2849–2861.e6, Nov. 2019. [Online]. Available: http://dx.doi.org/10.1016/j.celrep.2019.10.081

[58] Y.-C. Yu, R. S. Bultje, X. Wang, and S.-H. Shi, “Specific synapses develop preferentially among sister excitatory neurons in the neocortex,” Nature, vol. 458, no. 7237, p. 501–504, Feb. 2009. [Online]. Available: http://dx.doi.org/10.1038/nature07722

[59] J. A. Prasad, A. H. Balwani, E. C. Johnson, J. D. Miano, V. Sampathkumar, V. de Andrade, K. Fezzaa, M. Du, R. Vescovi, C. Jacobsen, and et al., “A three-dimensional thalamocortical dataset for characterizing brain heterogeneity,” biorXiv, May 2020. [Online]. Available: http://dx.doi.org/10.1101/2020.05.22.111617

[60] M. Han, H. Kim, G. Gu, K. Park, and W.-S. Han, “Efficient Subgraph Matching,” Proceedings of the 2019 International Conference on Management of Data - SIGMOD’19, 2019. [Online]. Available: http://dx.doi.org/10.1145/3299869.3319880

[61] V. Vassilevska and R. Williams, “Finding, minimizing, and counting weighted subgraphs,” Proceedings of the 41st annual ACM symposium on Symposium on theory of computing - STOC’09, 2009. [Online]. Available: http://dx.doi.org/10.1145/1536414.1536477

[62] P. Hell and J. Nešetřil, “Colouring, constraint satisfaction, and complexity,” Computer Science Review, vol. 2, no. 3, p. 143–163, Dec. 2008. [Online]. Available: http://dx.doi.org/10.1016/j.cosrev.2008.10.003

